# A Biomechanical Model of Tumor-induced Intracranial Pressure and Edema in Brain Tissue

**DOI:** 10.1101/480673

**Authors:** I. C. Sorribes, M. N. J. Moore, H. M. Byrne, H. V. Jain

## Abstract

Brain tumor growth and tumor-induced edema result in increased intracranial pressure (ICP), which, in turn, is responsible for conditions as benign as headaches and vomiting, or as severe as seizures, neurological damage, or even death. Therefore, it has been hypothesized that tracking ICP dynamics may offer improved prognostic potential in terms of early detection of brain cancer and better delimitation of the tumor boundary. However, translating such theory into clinical practice remains a challenge, in part, due to an incomplete understanding of how ICP correlates with tumor grade. Here, we propose a multiphase mixture model that describes the biomechanical response of healthy brain tissue – in terms of changes in ICP and edema – to a growing tumor. The model captures ICP dynamics within the diseased brain and accounts for the ability/inability of healthy tissue to compensate for this pressure. We propose parameter regimes that distinguish brain tumors by grade thereby providing critical insight into how ICP dynamics vary by severity of disease. In particular, we offer an explanation for clinically observed phenomena such as lack of symptoms in low grade glioma patients versus a rapid onset of symptoms in those with malignant tumors. Our model also takes into account the effects tumor-derived proteases may have on ICP levels and extent of tumor invasion. This work represents an important first step towards understanding the mechanisms that underlie the onset of edema and ICP in cancer-afflicted brains. Continued modeling effort in this direction has the potential to make an impact in the field of brain cancer diagnostics.

## 1 INTRODUCTION

Gliomas arise from non-neuronal brain cells such as neuroglia or glial cells and are the most common type of primary brain tumors diagnosed in the United States each year [1]. Per World Health Organization (WHO) guidelines [2], gliomas are classified by grade, which is based on histological and molecular assessments of tumor biopsy samples. In particular, grade I gliomas are slow growing, non-malignant, and associated with a better prognosis than grade IV gliomas, which are highly malignant and invasive, with a mean overall survival of 15 months post-diagnosis [1, 3]. Each year, 23,000 new cases of glioma are diagnosed in the US and it is responsible for over 16,000 deaths [4, 5]. This high mortality rate is, in part, due to relatively modest advances in therapies over the last 25 years [6, 7]. The standard course of treatment involves a combination of surgery, radiotherapy, and chemotherapy. This treatment remains palliative rather than curative due to multiple factors such as late detection of high grade tumors and an inability of current imaging techniques to accurately capture the tumor boundary, especially in the case when the cancer has infiltrated healthy tissue [6].

As brain cancers grow, tumor cells displace adjacent healthy tissue which, together with tumor-induced vascular abnormalities, causes a disruption of the blood-brain barrier. As a result, a large volume of blood plasma to leak into the tumor tissue causing cerebral edema, that is, abnormal swelling in the brain parenchyma [8]. Since the cranial vault is an enclosed and environmentally controlled space, a growing tumor – and any associated cerebral edema – may lead to an increase in intracranial pressure (ICP) thereby disrupting the homeostatic environment within the brain. It is this increase in ICP that is responsible for many adverse symptoms associated with brain cancers such as headaches, nausea, and seizures. Left unchecked, ICP may ultimately cause patient death [9]. For these reasons, it has been hypothesized that tracking the degree of edema or changes in ICP may offer improved prognostic potential in terms of early detection of disease and better delimitation of the tumor boundary [6]. This has motivated the recent development of new ultrasound techniques, such as shear wave elastography (SWE), which can detect changes in ICP that may be caused by a growing tumor [10].

Our goal in this paper is to explicate how edema and elevated ICP develop in brains afflicted with cancer. In order to realize this aim, we propose a mathematical model of the biomechanical effects of cancer on adjacent, healthy brain tissue. Specifically, a multiphase framework is employed to describe the physical forces and interactions that are involved in the response of brain tissue to a growing tumor. Using analytical and numerical techniques, we explore parameter regimes that correspond to different tumor grades and establish how these may correlate with ICP time-courses. Our model, if validated with experimental data, has the potential to accelerate the clinical applicability of new diagnostic modalities such as SWE. Indeed, mathematical models have been employed extensively to describe the growth and response to treatment of gliomas (for instance, see [11-16]; we refer the reader to [17] for a recent review). However, the focus of these previous models has been to predict either the extent to which the tumor has invaded brain tissue, or the outcome of different treatment strategies. To the best of our knowledge, ours is the first model to explicitly consider tumor-induced changes in ICP.

The remainder of this manuscript is organized as follows. In section 2, we present our mathematical model and describe the numerical scheme employed to simulate our model. We also provide a discussion on parameter values and constitutive assumptions. In section 3, we employ singular perturbation techniques to study parameter regimes that correspond to brain cancer by grade, and present numerical validation of our analysis. In section 4, we extend our model to investigate the effect of proteases released by the tumor on the emergence of peritumoral edema. Finally, we conclude with a discussion on the significance of our findings in section 5.

## 2 MATHEMATICAL MODEL

### 2.1 Model Derivation

A multiphase framework is used to describe the response of healthy brain tissue to a growing tumor. Specifically, brain tissue is viewed as a mixture of two distinct phases: a cellular phase comprising healthy tissue; and an aqueous phase consisting primarily of water, and dissolved extracellular tissue components. Mass and momentum balances are applied to both phases. The resulting equations are closed by imposing suitable constitutive relations for mass exchange between the different phases, the partial stress tensors, and momentum transfer between the phases. The mass exchange terms are defined on the basis of phenomenological observations of cell growth whereas the choice of stress tensors and momentum transfer terms are based on the assumed mechanical properties of each phase.

Indeed, multiphase models have been used extensively to simulate various aspects of cancer biology such as tumor growth, tumor-extracellular matrix interactions [18], cancer infiltration [19-22] and tumor encapsulation [23]. We refer the reader to [24] for a review of such models. We remark that although we follow the general framework provided in [20, 23], our model differs significantly from those preceding it in that we are focused on the biomechanical response of healthy tissue to a growing (brain) tumor, rather than the tumor itself, or the tumor’s response to therapeutic intervention.

For simplicity, we formulate our model in one-dimensional Cartesian geometry with *x* = 0 representing the skull and *x* = *G*(*t*) representing the boundary of a growing tumor. Specifically, the tumor is assumed to occupy the region *G*(*t*) < *x* < *L* and the tumor-healthy tissue boundary moves with a fixed velocity *v*, which is assumed to correlate with tumor growth rate. That is, *G*(*t*) = *G*(*t* = 0) − *vt* where *G*(*t* = 0) = *L*. We remark that, since our focus is on modeling the development of edema and ICP in the healthy tissue surrounding the tumor, we do not explicitly model tumor growth. A model schematic is shown in figure 1.

**Figure 1:**
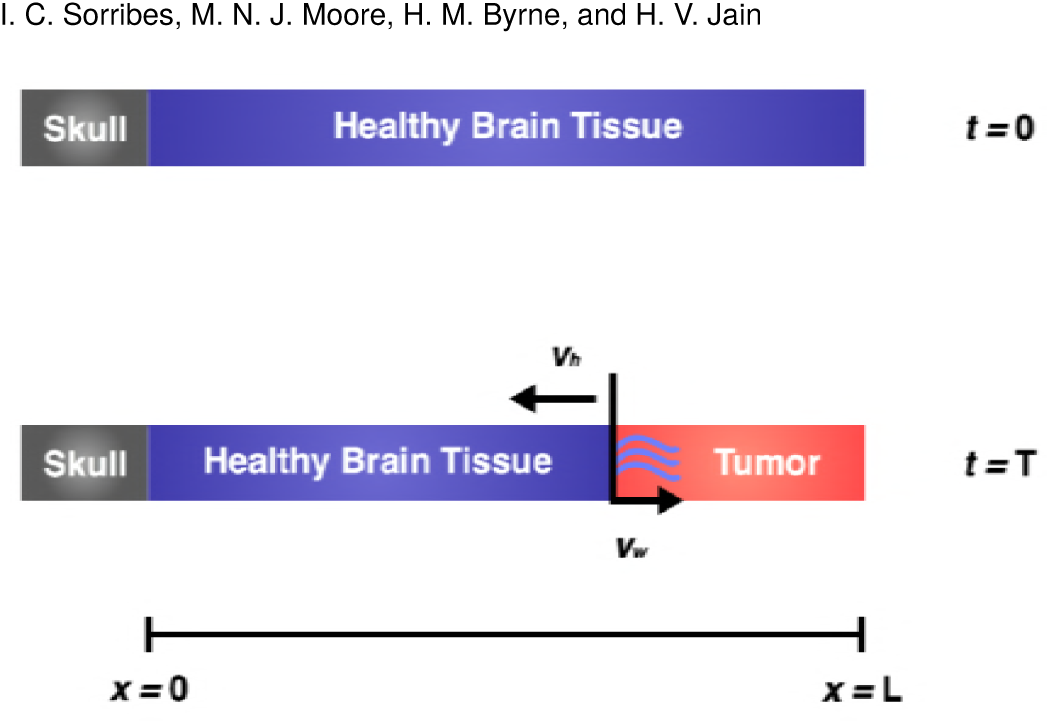
Schematic showing the geometry of the model domain. The skull is located at *x* = 0 and *x* = *G*(*t*) represents the boundary of a growing tumor, which is assumed to occupy the region *G*(*t*) < *x* < *L*, shown in red. The tumor-tissue interface is assumed to move into healthy tissue with a fixed velocity *v* that is representative of tumor growth. This boundary is permeable to fluid but impermeable to healthy tissue and the skull is impermeable to both, tissue and fluid.

We denote the volume fraction of healthy tissue by *h*(*x*, *t*) and that of the aqueous phase by *w*(*x*, *t*). Their corresponding velocities and stress tensors are denoted *v*_*h*_, *v*_*w*_, *σ*_*h*_, and *σ*_*w*_ respectively, these quantities being scalars for the one-dimensional geometry considered here. Both phases are assumed to be incompressible fluids whose densities are, to leading order, equal. In consequence, mass balances for each phase may be expressed as:

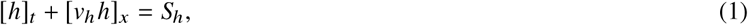

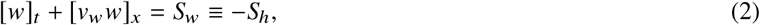

where [·]_*t*_ and [·]_*x*_ denote partial derivatives with respect to time and space, and *S*_*h*_ and *S*_*w*_ denote the net rates of production of healthy cells and water, respectively. We assume that the system is closed and, hence, there is no net volume change: the volume is simply transformed from one phase to another. Assuming further that there are no voids within the region, we obtain:

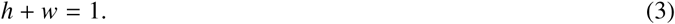

Following [23], healthy cells are assumed to proliferate by absorbing water at a rate which is proportional to *h* and *w*, and they undergo apoptosis at a rate which is proportional to *h*. Taken together, these assumptions yield:

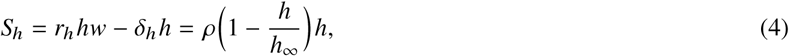

where *ρ* = (*r*_*h*_ - *δ*_*h*_) is the net growth rate of healthy cells and *h*_∞_ = (1 - *δ*_*h*_/*r*_*h*_ represents their steady-state volume fraction in the absence of external stimuli.

Next, neglecting inertial effects and assuming that no external forces act on the system, the momentum conservation laws may be written as follows:

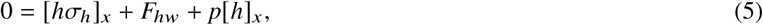

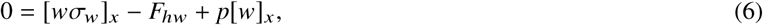

where *F*_*hw*_ is the force exerted on healthy tissue by the aqueous phase, an equal and opposite force being exerted by healthy tissue on the aqueous phase. We view the cells and aqueous fluid as inviscid fluids and, following [23, 25], we prescribe the stress tensors *σ*_*h*_, *σ*_*w*_ to be:

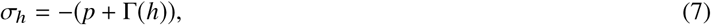

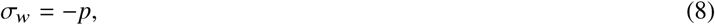

where *p* is the assumed pressure common to both phases and Γ (*h*) represents an additional isotropic pressure that distinguishes healthy tissue from water. Following [23], the interaction or drag term *F*_*hw*_ is taken to be proportional to the relative velocities of the phases, and to depend linearly on their volume fractions, so that:

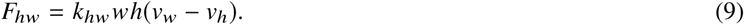

The equations governing the dependent variables *h*(*x*, *t*), w(*x*, *t*), *v*_*h*_, *v*_*w*_, and *p* are closed by prescribing Γ(*h*) and appropriate boundary and initial conditions. Specifically, brain tissue is taken to be at its spatially-homogeneous equilibrium state initially, that is, *h*(*x*, 0) = *h*_∞_. The skull is assumed to be impermeable to both tissue and water so that *v*_*h*_ = 0 = *v*_*w*_ at *x* = 0. Brain cancer growth is known to disrupt the blood-brain barrier [8]; therefore we assume that the tumor is permeable to fluid but impermeable to the solid tissue phase. That is, *v*_*h*_ = -*v* at *x* = *G*(*t*) where *G*(*t*) = *L* -*vt*. In practice, water may flow from the tumor into the healthy tissue or vice versa. Here we assume that once fluid enters the tumor it cannot escape.

### 2.2 Model Simplification

We now reduce the two-phase model developed above to a single nonlinear parabolic equation for the volume fraction of the cellular phase. We eliminate *w* = 1 - *h* from the model equations via the “no voids” assumption in (3). Then, adding (1) and (2):

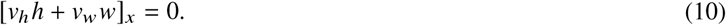

Integrating the above with respect to *x* and recalling that the skull (at *x* = 0) is impermeable to fluid and tissue, we obtain:

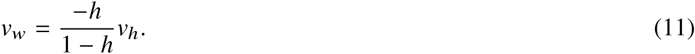

Adding (5) and (6), and using (3) and (7)-(8), the momentum balance for the system reduces to:

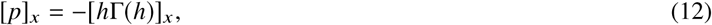

that is *p*(*x*, *t*) = -*h*Γ(*h*) + *p*_0_(*t*). We now substitute from (4),(8),(9), (11), and (12) into (6) to obtain the following expression for the velocity of the tissue phase:

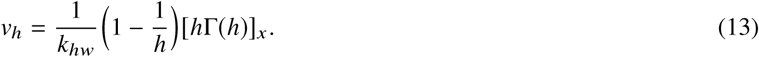

We note that the positive constant *k*_*hw*_ may be absorbed into Γ(*h*) and therefore we neglect it in what follows. Finally, we substitute from (13) into (1), to arrive at the following partial differential equation (PDE) describing the biomechanical response of healthy tissue to a growing tumor:

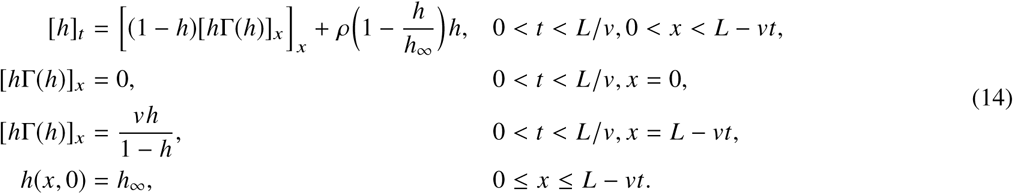

### 2.3 Numerical Methods

Our challenge now is to numerically solve the nonlinear reaction-diffusion system (14) with a moving boundary. We first transform the moving domain [0, 1 - *vt*] to a fixed one [0,1] via:

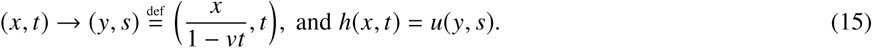

Consequently, (14) transforms to:

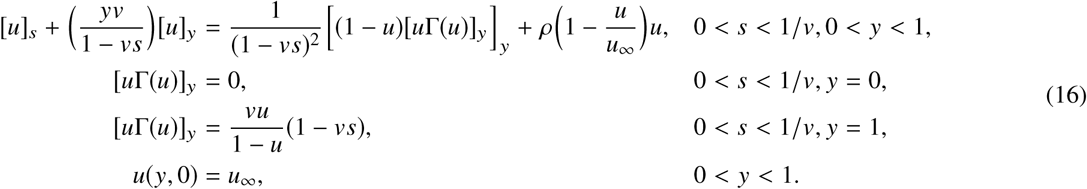

Note that the coordinate transformation introduces an advection term on the left-hand-side of the PDE. Thus, (16) is an advection-reaction-diffusion PDE with variable-coefficient advection and nonlinear diffusion. To integrate this system forward in time, we employ operator splitting, which has the advantage of superior stability [26]. In particular, we integrate the reaction-diffusion term over one time step using Crank-Nicolson, with the nonlinearities lagged. This provisional solution is then advected using upwinding to obtain *u* at the next time step. Note that the advection term vanishes at the left boundary which allows the associated boundary condition to be safely omitted from the advection step. The overall accuracy of this scheme is second-order in space and first-order in time. Further details on the numerical scheme and its validation with a test problem can be found in the Supplemental Information.

### 2.4 Parameter Values and Functional Forms

Equations (14) are completely determined by appropriate choices of the volume fraction *h*_∞_ of healthy brain at homeostasis, the speed *v* at which the tumor-healthy tissue boundary moves, the rate *ρ* of healthy brain tissue remodeling, and the additional isotropic pressure Γ (*h*) in the tissue fraction. Without loss of generality, we set *h*_∞_ = 0.5. The choice of *v* and *ρ* will determine the type of brain cancer being simulated. Therefore, in section 3, we investigate how the system dynamics change as *v* and *ρ* are varied.

Finally, we discuss our choice for Γ(*h*). Patients with slow growing tumors are initially asymptomatic due to minimal changes in their ICP. If the tumor continues to grow in size, eventually different compensatory mechanisms of the brain will be exhausted, resulting in a sharp increase in ICP [27]. Langfitt et al. [28] proposed a qualitative relation between ICP and intracranial volume, shown in figure 2(a), that represents this sequence of disease progression. We choose a functional form for Γ(*h*) that reproduces the Langfitt curve in the case when *v* is small compared to other model parameters. Specifically, we take:

**Figure 2:**
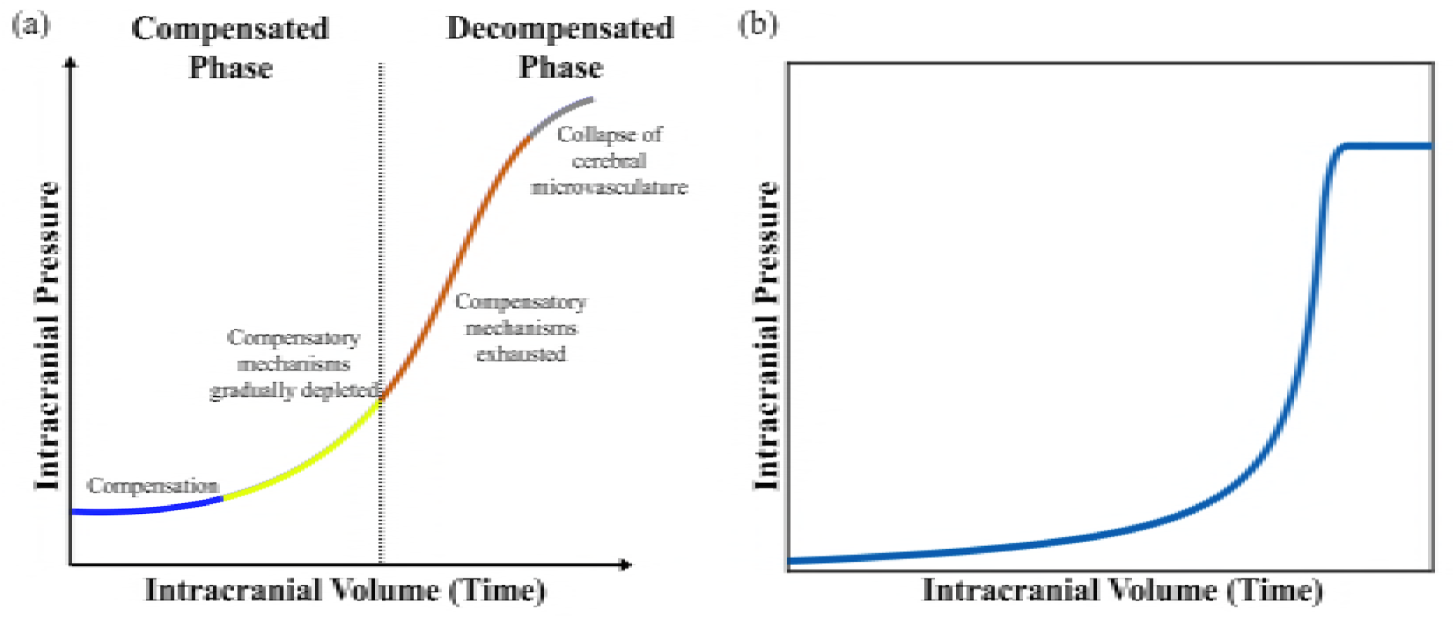
(a) Reproduction of the qualitative Langfitt Curve following Figure 2 in [29] and Figure 2 in [30]. As intracranial volume increases with time, due to a growing tumor within the cranial vault, the compensatory mechanisms that avoid an initial rise in ICP are exhausted. This exhaustion results in the transition between a compensated phase and a decompensated phase, which eventually leads to a collapse of cerebral microvasculature. (b) Model output of average ICP versus time for simulations of equations (14) with *v* = 0.01 and *ρ* = 0.1.

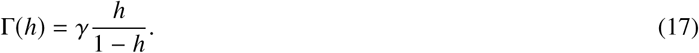

Figure 2(b) shows model predictions of the average ICP versus time for a specified set of model parameters (*γ* = 1, *v* = 0.01, *ρ* = 0.1), and it is observed to be in good qualitative agreement with the behavior of ICP as expected from the Langfitt curve. We calculated average ICP by integrating equation (12) and averaging the resulting pressure *p*(*x*, *t*) over space. We remark that other functional forms of Γ(*h*) are also possible; our choice was governed by reasons of simplicity and to keep the number of unknown parameters at a minimum.

To illustrate model dynamics for this choice of Γ(*h*), we present numerical simulations as model parameters are varied. Specifically, we set *γ* = 1 and vary *v* and *ρ*. Figure 3(a) shows the initial distribution of *h*(*x*, 0) versus *x*. Figures 3(b)-(e) show plots of *h*(*x*, *t*) versus *x* as the rate *ρ* of tissue remodeling is increased. For each value of *ρ*, different rates *v* of tumor growth are considered. The simulations are run until *t* = *t*_*f*_ where *t*_*f*_ is the time when either *h*(*x*, *t*) approaches one, (at which point *v*_*w*_ → ∞, see equation (11)), or the tumor boundary *G*(*t*) reaches *x* = 0, whichever happens first. The plots are snapshots of the volume fraction of healthy tissue at this final time. As the tumor grows, water leaks into the tumor space and consequently, the volume fraction of healthy tissue increases across the simulation domain. Since we do not model tumor growth explicitly, we cannot quantify fluid accumulation or edema in the tumoral space. Rather, any increase in *h* above its equilibrium value *h*_∞_ is taken to be representative of peritumoral edema. In all cases considered, when *v* is small compared to *γ*, (dashed red and dotted blue curves) the healthy tissue fraction appears to increase uniformly across the domain as the tumor grows. This is because the diffusion of healthy tissue over space overcomes the pushing effect of the tumor boundary. As a consequence, the tumor can invade far into the healthy tissue. Moreover, when *ρ* is large and *v* is small, (figure 3(d), dashed red and dotted blue curves), remodeling ensures that the healthy tissue fractions stay below one longer so that the tumor invades the furthest. This observation is summarized in figure 3(f), which shows the maximum degree of tumor invasion into healthy tissue as *v* and *ρ* are varied. On the other hand, when *v* is comparable to *γ*, *h*(*x*, *t*) →1 rapidly at the tumor boundary and the tumor is predicted to cease growth due to a lack of space. Biologically, this could mean that the primary tumor mass has reached a critical size. At this point, individual tumor cells may be shed from the primary tumor and invade into healthy tissue, as is known to happen in glioblastomas [31].

**Figure 3:**
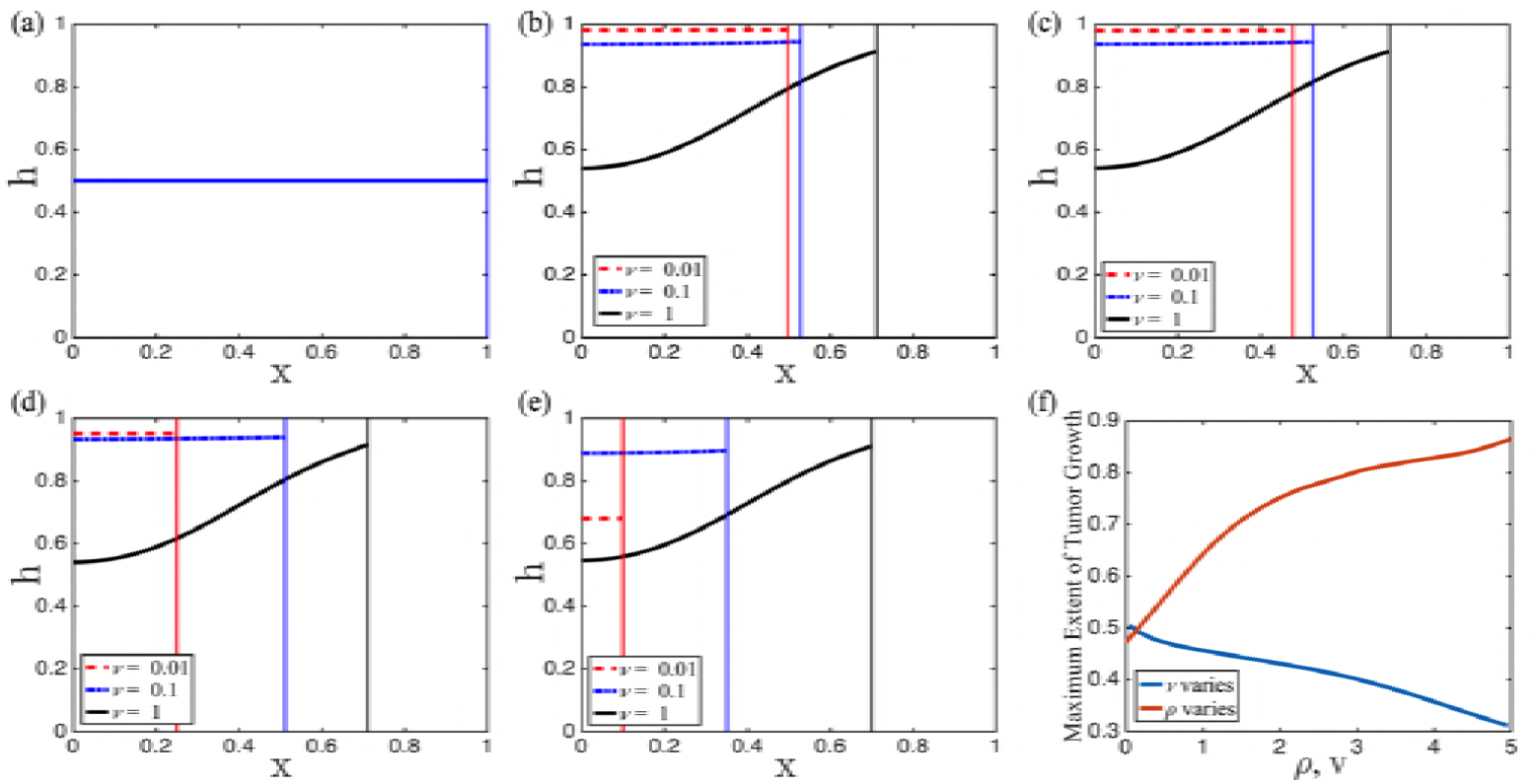
(a) Initial distribution healthy tissue *h*(*x*, *t*) versus *x*. (b)-(e)Plots of *h*(*x*, *t*) versus *x* as the rate *ρ* of tissue remodeling is increased ((b), *ρ* = 0, (c), *ρ* = 0.01, (d), *ρ* = 0.1, and (e), *ρ* = 1). In each case, different rates *v* of tumor growth are considered (dashed red curve: *v* = 0.01, dash-dotted blue curve: *v* = 0.1, and solid black curve: *v* = 1). Simulations of (14) were run until either *h*(*x*, *t*) approaches one, or the tumor boundary reaches *x* = 0. The vertical lines denote the position of the tumor. (f) Distance invaded by the tumor into healthy tissue as *v* and *ρ* are varied with *γ* = 1.

We also present a heat map of pressure corresponding to some of the cases discussed above. Specifically, we choose *ρ* = 0 (figure 4, top row) and *ρ* = 0.1 (figure 4, bottom row) for illustrative purposes. For each value of *ρ*, we consider a slow, medium and fast growing tumor, as before. These pressure plots correspond to the profiles in figures 3(b) and (d) respectively. Time is plotted on the *y*-axis, and space on the *x*-axis. For ease of visualization, the pressure plots are shown in the transformed stationary domain, with the skull at *x* = 0, and the tumor boundary at *x* = 1. In all cases, when *v* is small compared to *γ* (figures 4(a),(b),(d),(e)), ICP is predicted to increase almost uniformly across the domain with minimal increase initially when the tumor is small. Such a patient may remain asymptomatic during the initial stages of tumor growth, with symptoms manifesting only when ICP changes appreciably. For a faster growing tumor (figures 4(c),(f)) pressure throughout the domain barely changes except near the tumor. These predictions reflect what is observed clinically, that is, sharp increases in ICP when compensatory mechanisms are exhausted [27, 32, 33], lending credibility to our model.

**Figure 4:**
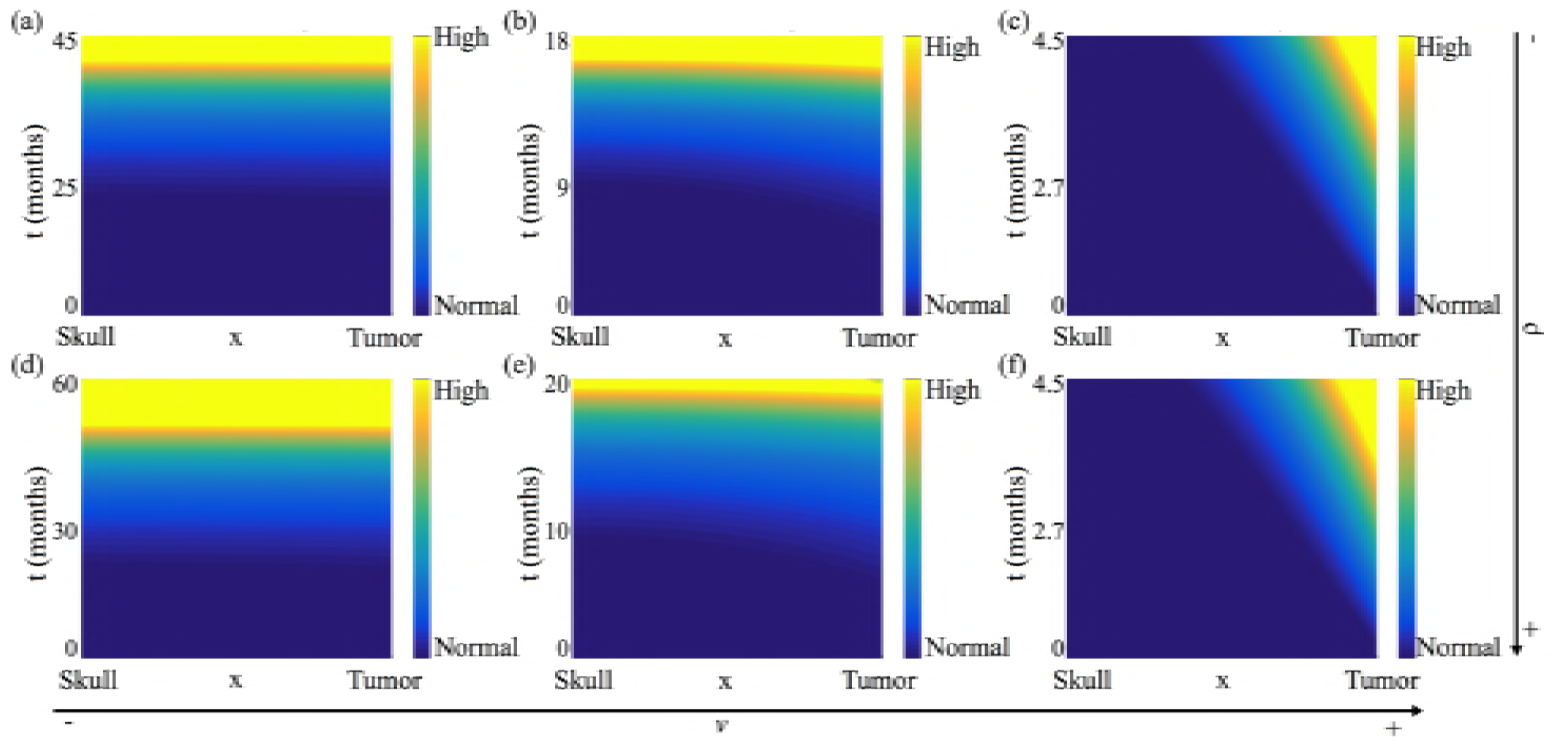
Heat map of pressure corresponding to the healthy tissue profiles shown in figures 3(b),(d). Time, measured in months, is plotted on the *y*-axis and space on the *x*-axis. For ease of visualization, the pressure plots are shown in the stationary domain with the skull located at the minimum of the *x*-axis, and the tumor boundary at the maximum of the *x*-axis. (a)-(c) correspond to *ρ* = 0 and (d)-(f) correspond to *ρ* = 0.1. For each value of *ρ*, different rates *v* of tumor growth are considered ((a), (d), *v* = 0.01, (b), (e), *v* = 0.1, and (c), (f), *v* = 1).

Finally, for illustrative purposes, we also present numerical simulations of the case when Γ(*h*) is a constant rather than an increasing function of *h*(*x*, *t*). Figure 5 presents plots of *h*(*x*, *t*) versus *x* as *ρ* is increased, with different tumor growth rates *v* considered for each value of *ρ* as before. The corresponding pressure plots are shown in figure 6. Qualitatively, figures 3 and 5 are similar; they both predict that slow growing tumors can invade farther into the brain, but for faster growing tumors, the primary tumor mass ceases growth sooner. Thus, these graphs are not by themselves indicative of whether Γ(*h*) = *γ* or Γ(*h*) = *γ*(*h*)/(1 – *h*) is a more biologically appropriate functional form for the additional isotropic pressure in the tissue phase. However, when we look at predicted changes in ICP shown in figure 6, we observed that when *v* is small and Γ(*h*) is constant, ICP increases linearly and does not match the Langfitt curve.

**Figure 5:**
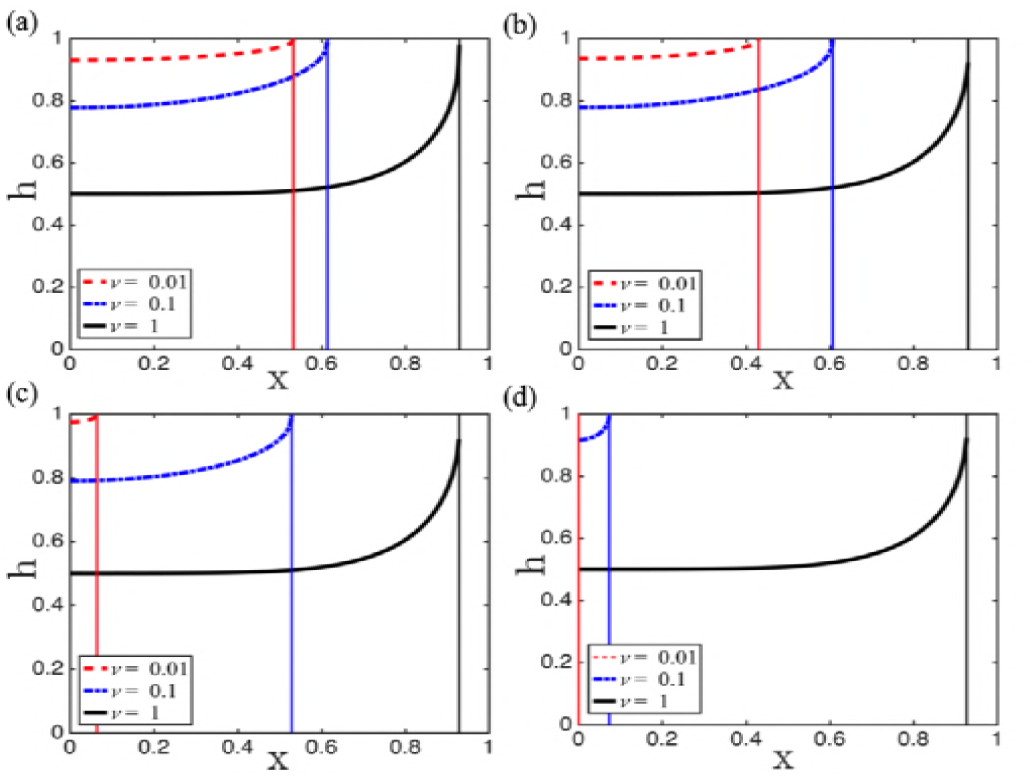
(a)-(d) Plots of *h*(*x*, *t*) versus *x* when the additional isotropic pressure in the tissue phase is constant, that is, Γ(*h*) = *γ*, and the rate *ρ* of tissue remodeling is increased ((a), *ρ* = 0, (b), *ρ* = 0.01, (c), *ρ* = 0.1, and (d), *ρ* = 1). In each case, different rates *v* of tumor growth are considered (dashed red curve: *v* = 0.01, dash-dotted blue curve: *v* = 0.1, and solid black curve: *v* = 1). Simulations of (14) were run until either *h*(*x*, *t*) approaches one, or the tumor boundary reaches *x* = 0. The vertical lines denote the position of the tumor.

**Figure 6:**
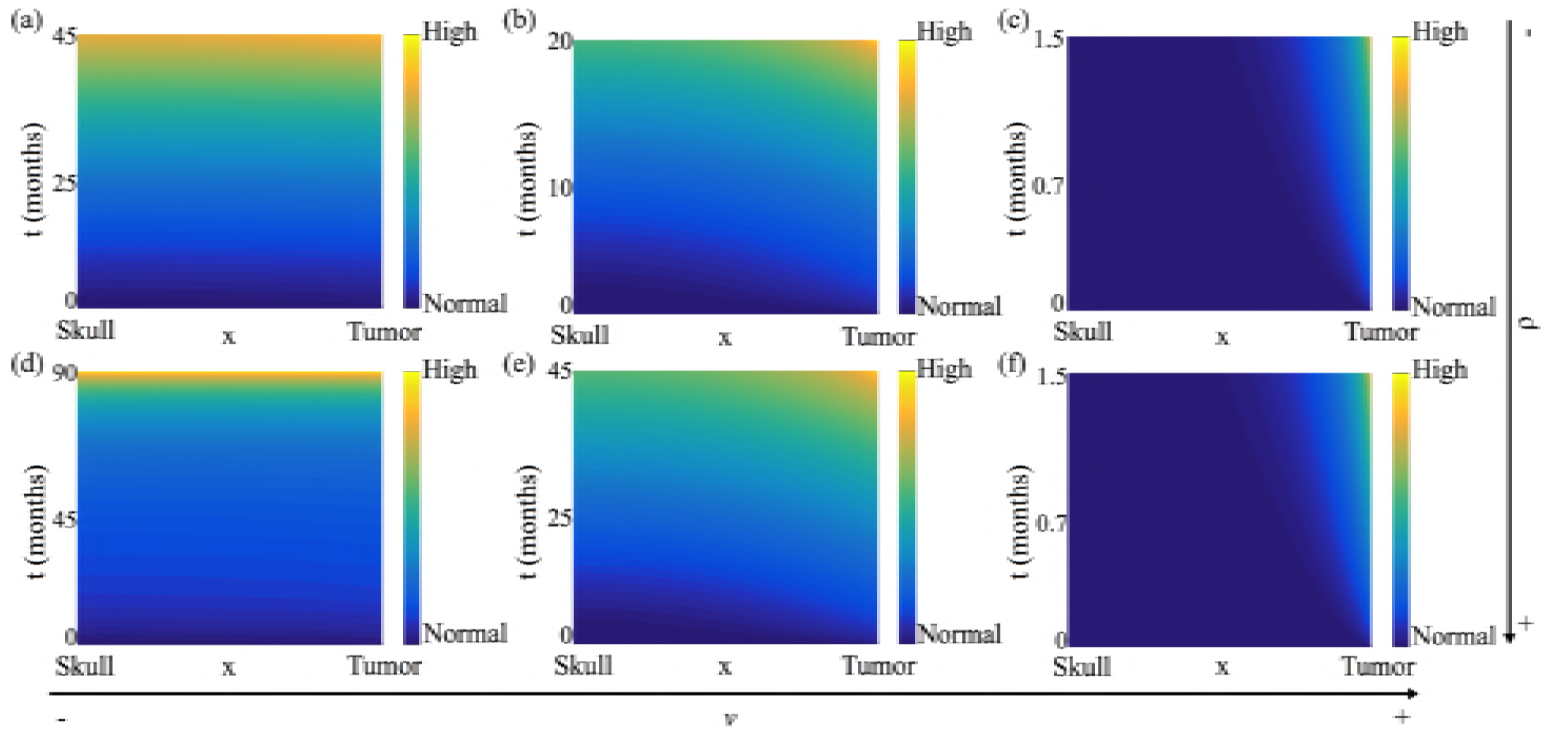
Heat map of pressure corresponding to the healthy tissue profiles shown in figures 5(a),(c) when the additional isotropic pressure in the tissue phase is constant, that is, Γ(*h*) = *γ*. For ease of visualization, the pressure plots are shown in the stationary domain with the skull and the tumor boundary located at the minimum and maximum of the *x*-axis, respectively. Time, measured in months, is plotted on the *y*-axis. (a)-(c) correspond to *ρ* = 0 and (d)-(f)correspond to *ρ* = 0.1. For each value of *ρ*, different rates *v* of tumor growth are considered ((a), (d), *v* = 0.01, (b), (e), *v* = 0.1, and (c), (f), *v* = 1.

## 3 ASTROCYTOMAS BY GRADE: SCALING REGIMES AND SIMULATIONS

As mentioned in the introduction, brain tumors are classified by grade I to IV, where grade I is a benign tumor and grade IV the most malignant [2]. In our model *γ* (additional isotropic pressure), *ρ* (healthy tissue remodeling), and *v* (tumor growth rate) are critical determinants of the response of healthy tissue to a growing tumor in terms of: changes in ICP; how far the tumor can invade into healthy tissue; and peritumoral edema, or change in tissue volume fraction above a normal state. We now propose regimes of parameter space that correspond to each grade, focusing on astrocytomas, the most common and malignant type of gliomas [1, 34]. However, similar analyses may be conducted for other types of brain cancer.

Specifically, we identify and analyze distinguished limits of the parameters *γ*, *v* and *ρ* that yield distinct profiles for ICP, the degree of tumor invasion and deviation of tissue volume fraction from its healthy steady state. These profiles are then compared qualitatively to clinical observations, and supported by numerical simulations for typical cases. We restrict our attention to *γ* = 𝒪(1) and *γ* = 𝒪 (1/*∈*). Note that *γ* = 𝒪(*∈*) is biologically unrealistic since in such a regime there will be no increase in ICP as the tumor grows. In particular, we focus on *γ* = 𝒪(1/∈); *γ* = 𝒪(1) yields qualitatively similar results. Time is rescaled, if necessary, so that significant tumor growth can be observed.

### 3.1 Pilocytic Astrocytoma (grade I)

Pilocytic astrocytomas are benign, slow growing tumors of moderate cellularity. Patients with grade I astrocytomas have a good prognosis in general because gross total resection is typically curative [35]. Despite this, these tumors can become very large and remain asymptomatic for prolonged periods of time due to minimal changes in ICP [33, 36].

In our model, taking *v* = 𝒪(*∈*) and *ρ* = 𝒪(*∈*^2^) corresponds to a pilocytic astrocytoma. This can be seen by performing the transformation 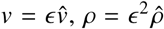and 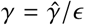in equations (14). Additionally, we rescale time to be on a long timescale *t* ∼ 𝒪(1/*∈*) so that we can observe significant tumor growth, that is, we define τ = *∈t*. Then, equations (14) reduce to:

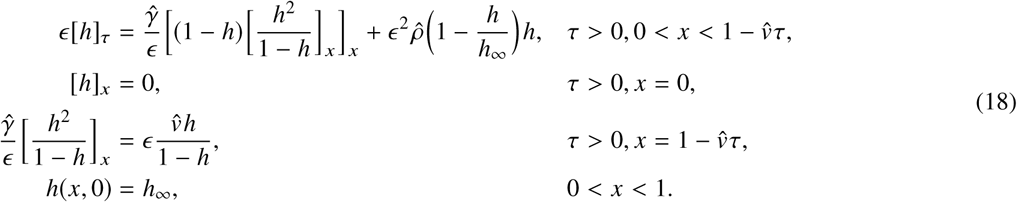

We seek solutions of equations (18) in the form of a regular power series expansion, with *h* = *h*_0_ + *∈h*_1_ + 𝒪(*∈*^2^). At leading order, we obtain:

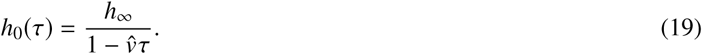

Thus, we predict that the volume fraction of the healthy tissue will increase uniformly across the simulation domain as the tumor boundary moves. Higher order correction terms may be determined in a similar manner (details of this analysis are presented in **??** of the Supplemental Information, together with figures showing a comparison of *h*(*x*, *t*) time-courses for the full model and the asymptotic solution). Figure 7(a) shows a heat map of the predicted pressure under this parameter regime. As expected, the slowly growing tumor invades the healthy tissue without a significant build-up of ICP. Our results are therefore consistent with clinical observations since grade I astrocytomas are benign and remain asymptomatic for a long time [33, 36].

**Figure 7:**
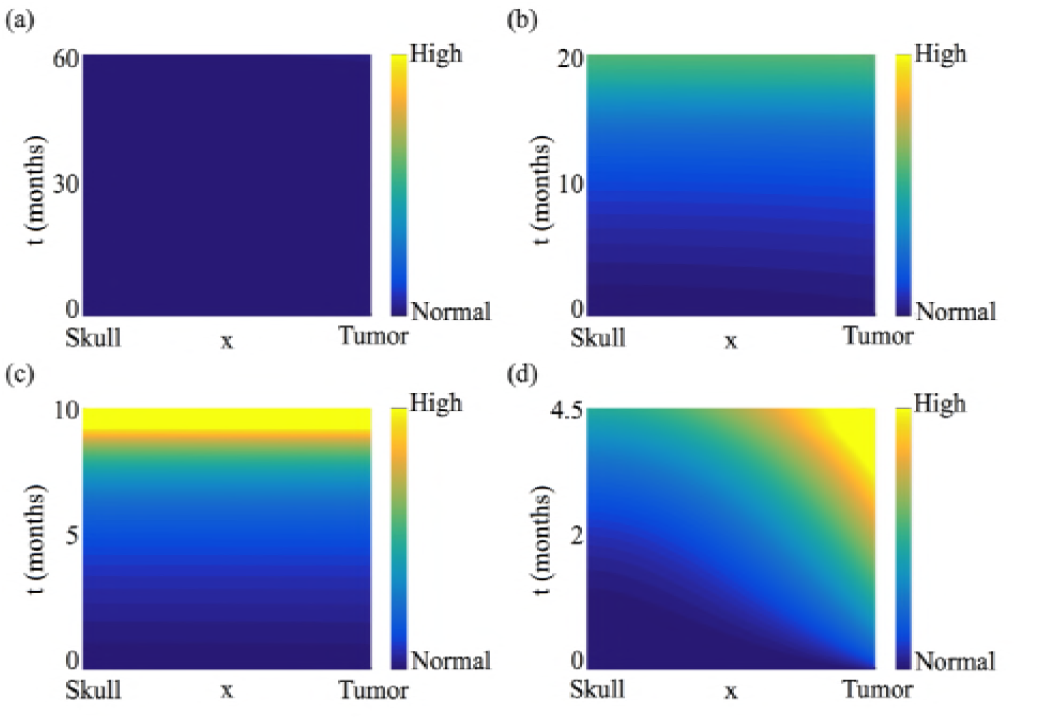
Pressure heat maps for parameter regimes corresponding to tumors by grade: (a) Grade I, Pilocytic Astrocytoma, *ρ* = 𝒪(*∈*^2^), *v* = 𝒪(*∈*); (b) Grade II, Diffusive Astrocytoma, *ρ* = 𝒪(1), *v* = 𝒪(1); (c) Grade III, Anaplastic Astrocytoma, *ρ* = 0, *v* = 𝒪(1); and (d) Grade IV, Glioblastoma, *ρ* = 𝒪 (*∈*), *v* = 𝒪(1/*∈*). In all cases, *γ* = 𝒪(1/*∈*).

### 3.2 Diffusive Astrocytoma (grade II)

Diffusive astrocytomas grow faster than pilocytic astrocytomas but slower than tumors of grades III and IV, and have ill-defined boundaries. Consequently, they cannot easily be cured by surgical resection alone. Patients with diffusive astrocytomas typically survive for 5 to 8 years and display symptoms related to increased ICP [33, 35, 36].

We show below that taking *v* = 𝒪(1) and *ρ* = 𝒪(1) corresponds to a diffusive astrocytoma. In this case remodeling and tumor growth occur on the same timescale and this is shorter than that for the grade I pilocytic astrocytoma considered in section 3.1. Additionally, with 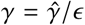(as before), equations (14) supply:

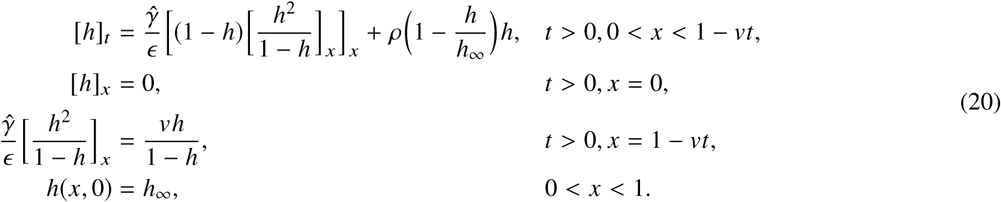

As before, we look for regular power series expansions of the form *h* = *h*_0_ + *∈h*_1_ + 𝒪(*∈*^2^). It is straightforward to show that the leading order term is given by:

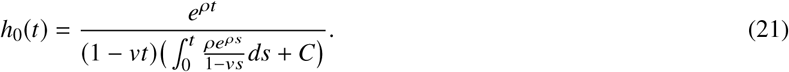

Higher order correction terms may be determined in a similar manner. Figures comparing time-courses of *h*(*x*, *t*) for the full model and the asymptotic solutions can be found in the Supplemental Information (Figure **??**). Figure 7(b) shows a heat map of the pressure for this parameter regime. As expected, there is a slow but noticeable build-up of ICP, and the distribution of *h*(*x*, *t*) across the spatial domain is no longer uniform. Our results are consistent with clinical observations of diffusive astrocytomas which are relatively slow growing, but cause ICP to increase.

### 3.3 Anaplastic Astrocytoma (grade III)

Anaplastic astrocytomas are malignant tumors with a mean survival of three years [35]. They are more aggressive than grade I and II astrocytomas and tend to have tentacle-like projections [36]. In contrast to the previous cases, where the brain can adapt, to some extent, to the growing tumor, now the increase in ICP is more rapid. Consequently, symptoms for grade III (and also grade IV) astrocytomas appear more rapidly [33].

For anaplastic astrocytomas we assume that the tumor grows at a comparable speed as in the previous case, so that *v* = 𝒪(1). Since, the brain does not adapt as quickly to volume changes induced by tumor growth we fix *ρ* = 𝒪(*∈*). In particular we take, *ρ* = 0 for analytical tractability. Then, equations (14) reduce to:

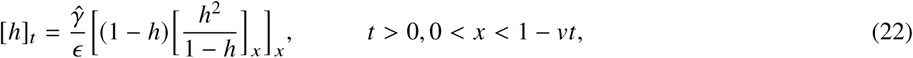

with initial and boundary conditions remaining unchanged from the previous case (grade II). By seeking solutions of the form *h* = *h*_0_ + *∈h*_1_ + 𝒪 (*∈*^2^), it is straightforward to show that:

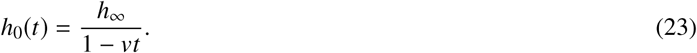

Thus, at leading order, anaplastic astrocytomas behave like pilocytic astrocytomas. However, the first correction term for anaplastic astrocytomas reveals spatial inhomogeneity. That is:

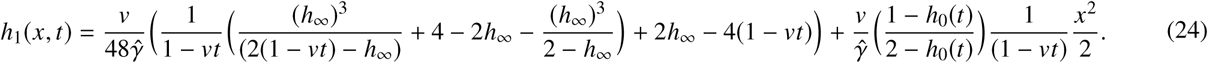

This is in contrast to the case of pilocytic astrocytomas where the first correction term is also independent of space. Details of this analysis are presented in **??** of the Supplemental Information, together with figures comparing time-courses for the full model and the asymptotic approximation (figure **??**). Figure 7(c) shows a heat map of the predicted pressure under this parameter regime. Our results are consistent with clinical observations in that the build-up of ICP is more rapid as compared to that in astrocytomas of grades I and II.

### 3.4 Glioblastoma Multiforme (grade IV)

Glioblastomas (GBM) are the most rapidly growing and malignant of astrocytomas [1]. GBM are extremely aggressive, and patients have an average survival period of 12 to 18 months post-diagnosis [35].

We show below that taking *v* = 𝒪 (1/ *∈*), so that 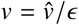, corresponds to GBM. As in the case of anaplastic astrocytoma, we assume *ρ* = 𝒪 (*∈*), that is, 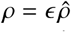, since the brain does not adapt quickly to volume changes induced by tumor growth. When modeling GBM we rescale time by setting *τ* = *t* /*∈* so that we can capture effects that act on the short, growth time scale. Under these assumptions, equations (14) reduce to:

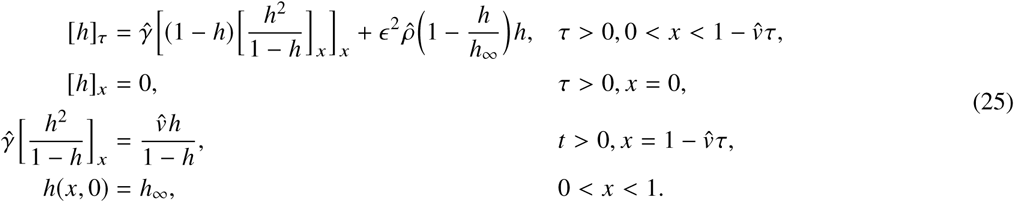

In this case, if we seek a regular power series expansion for *h*(*x*, τ), then we recover equations (14) at leading order. Therefore this case needs to be considered numerically. However, the asymptotic approximation for grade III cancer as given by (23) and (24) can be rescaled to approximate the initial stage of GBM. That is, introducing the transformation 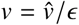 and *t* = *∈*τ in equations (23) and (24) supplies:

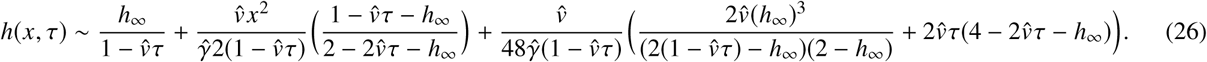

Figures comparing *h*(*x*, *t*) time-courses for numerical solutions of the full model and approximation (26) are included in the Supplemental Information (see figure **??**). Figure 7(d) shows a heat map of the pressure under this parameter regime. In agreement with clinical observations, our model simulations predict a rapid increase in ICP due to GBM growth. In contrast to the case for anaplastic astrocytomas, the increase in ICP is no longer spatially uniform, with the tumor-healthy tissue interface corresponding to the region of highest pressure.

## 4 APPLICATION: MODELING EDEMA WITH PROTEASES

The results presented in the previous sections reveal that our model captures key aspects of the mechanical response of brain tissue to tumor growth, and that it can be used to identify parameter regimes that correspond to tumor grades. One limitation of the model is that if the tumor growth rate *v* is large then all the fluid adjacent to the tumor leaks into it, causing the volume fraction of healthy tissue rapidly to approach one near the tumor boundary, which is biologically unrealistic. In practice, the mechanisms of tumor invasion are complex and depend, in part, on the secretion of several proteases that degrade healthy tissue and create space locally for tumor cells to invade and migrate [37]. In this section, we explain how we can extend our model to investigate the effect that tumor-derived proteases may have on ICP levels and tumor invasion.

Existing models [23, 38] typically view proteases as diffusible species. However, others argue that proteases act locally, in the immediate vicinity of the tumor source [39]. This point of view has been validated in recent experiments wherein protease expression was evaluated *in vivo* in a rat brain gliosarcoma xenograft model using magnetic resonance imaging. The experiments showed that proteases were localized in the peritumoral region [40]. Consequently, we assume that tumor-derived proteases will degrade healthy tissue adjacent to the moving tumor boundary, within a small region of width ∼ 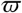. To account for this effect equations (14) are amended as follows:

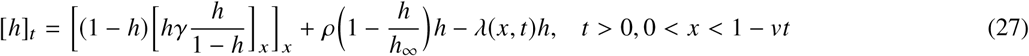

where the rate of tissue degradation λ(*x*, *t*) is assumed to be of the form:

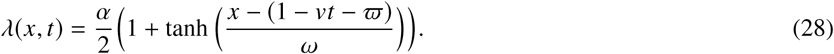

The initial and boundary conditions used to close equation (27) remain unchanged. In equation (28) the constant α represents the maximum rate at which proteases degrade healthy tissue and the constant *ω* determines the sharpness of the boundary of the region within which proteases are active. For illustrative purposes, we simulate the response of brain tissue to a fast growing tumor, such as GBM, and a slow growing tumor. Results for other malignancies can be similarly simulated. Figures 8(a), (b), and (c) show time snapshots of GBM growth, with and without active proteases. Specifically, figure 8(a) shows an early stage of tumor growth, figure 8(b) shows a snapshot at the time when the tumor not secreting proteases has invaded its furthest, and figure 8(c) shows the simulations at the time when the tumor secreting proteases has invaded its furthest. As can be seen, secreting proteases allows the tumor to keep growing for longer, with greater invasiveness. figure 8(d) and (e) graph the heat maps of corresponding changes in ICP with and without the effect of proteases, respectively. In both cases, a non-uniform increase in ICP is predicted as in the case of GBM earlier. However, ICP onset is slightly delayed with active proteases since the degradation of healthy tissue relieves the initial increase in ICP. Clinically this could explain why symptoms manifest late.

**Figure 8:**
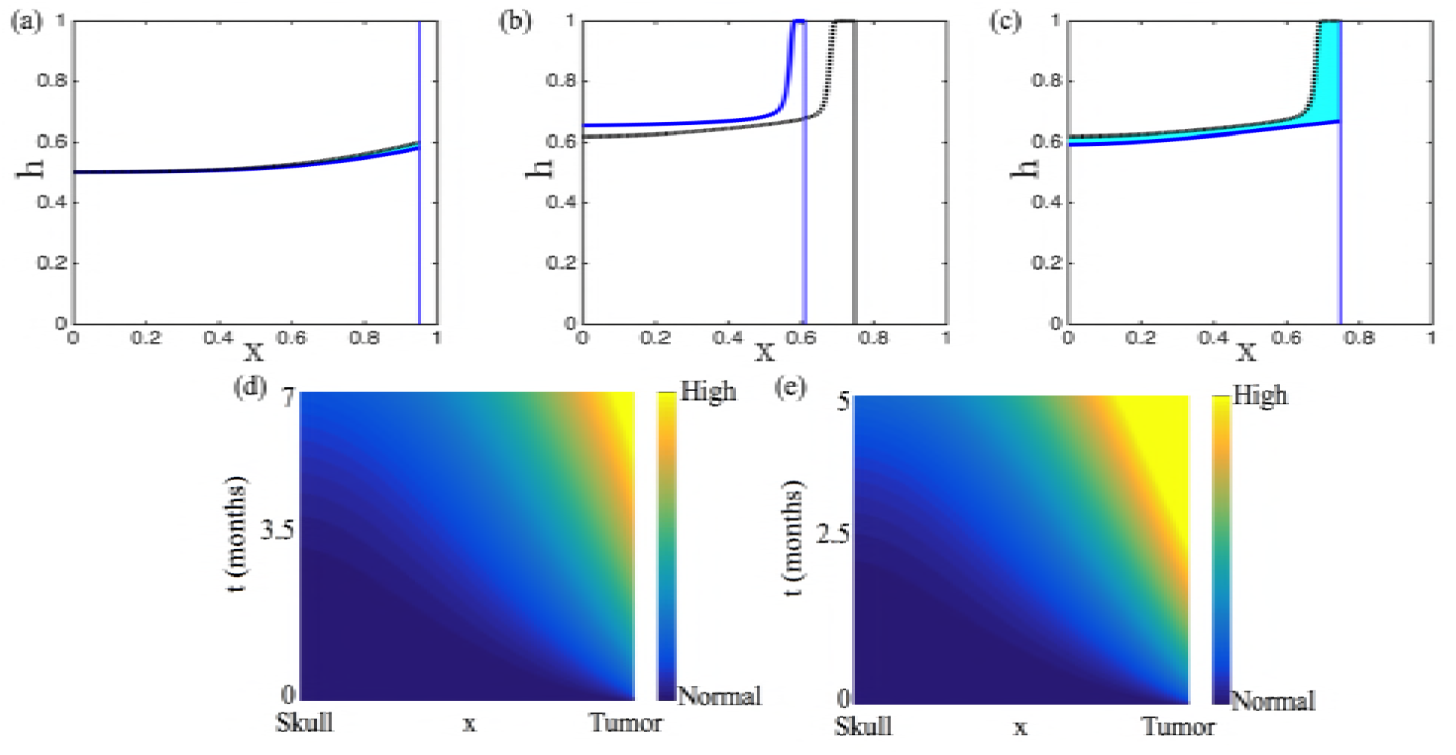
Numerical solution of (27) is presented in a blue solid line along with the solution of the equation without proteases (14) in a dashed black line. The area in between in shaded in cyan to illustrate the created edema. We consider parameters in the regime of a GBM (*γ* = 10, *ρ* = 0.01, *v* = 10, and *α* = 20). We present snapshots at an early time (a), the time when *h*(*x*, *t*) reaches 1 in GBM without proteases (b), and final times (c). Corresponding heat maps of pressure in GBM with (d) and without (e) the effect of proteases.

On the other hand, in figure 9(a), and (b) we present *h*(*x*, *t*) time snapshots for a slow growing tumor, with and without the effect of proteases. In this case, the tumor invades the simulation domain regardless of whether active proteases are present. However, the release of proteases results in a fluid build-up near the tumor, which we interpret as a component of peritumoral edema. As for pilocytic astrocytomas, there is minimal change in ICP regardless of whether tumor-derived proteases are present (figure 9(c)). These minimal changes in ICP are insignificant compared to the changes in ICP in the case of GBM.

**Figure 9:**
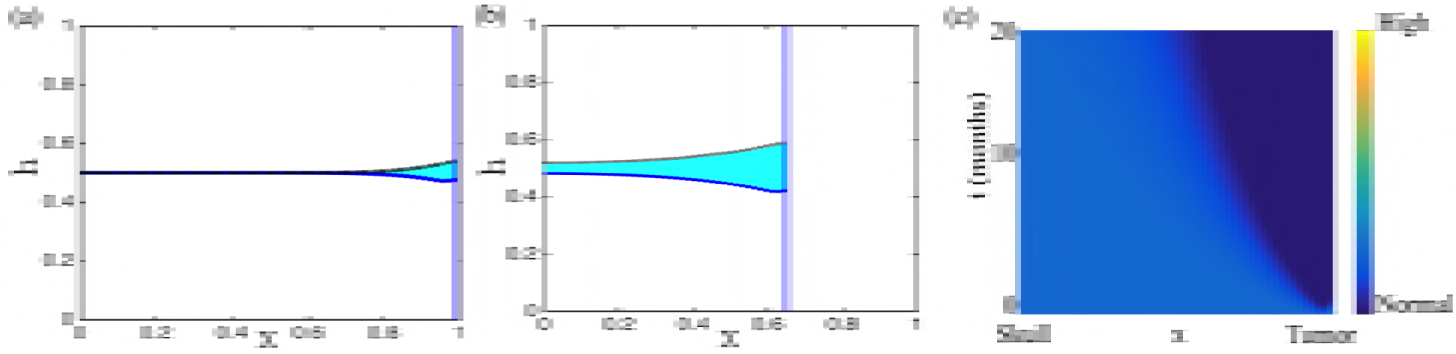
Numerical solution of (27) is presented in a blue solid line along with the solution of the equation without proteases (14) in a dashed black line. The area in between in shaded in cyan to illustrate the created edema. We consider the parameter regime of a slow growing tumor with *γ* = 0.01, *ρ* = 0.2, *v* = 0.01, and *α* = 0.4. We present snapshots at an early (a), and intermediate time (b). (c) Corresponding heat maps of pressure in a slow growing tumor with the effect of proteases.

## 5 DISCUSSION

Brain tumor growth disrupts the blood-brain barrier by displacing adjacent healthy tissue and inducing vascular abnormalities, and thereby causes cerebral edema [8, 9, 33, 35]. Since the brain is enclosed in the cranial vault, tumor growth and/or edema may cause an increase in ICP that manifests as headaches, nausea, and/or vomiting. Initially, these symptoms are subtle because of compensatory mechanisms such as cerebrospinal fluid displacement, cerebral blood flow decrease, and changes in the parenchyma shape. Over time, as the tumor progresses, such compensatory mechanisms are exhausted, and ICP increases sharply leading, eventually, to the patient’s death [27]. Currently, there is no clearly defined ICP threshold signaling the need for immediate treatment [41]. Understanding the relationship between ICP and tumor volume is crucial for determining such a threshold, and could be used as a new prognostic indicator.

Here, we proposed a biomechanical model of the response of brain tissue to a growing tumor in order to understand how factors such as tumor growth rate, healthy tissue remodeling rate, and the mechanical properties of brain tissue affect ICP. We viewed the brain as a two-phase mixture: a healthy tissue phase and a watery phase. The model was derived using principles of mass and momentum balances, and numerically integrated using an operator splitting scheme. Tumor growth was simulated by shrinking the right boundary of the domain. Model simulations and analysis provided critical insight into how edema and ICP depend on brain tumor by grade. In particular, we proposed parameter regimes that capture the differences in ICP dynamics associated with different grades of astrocytomas and used perturbation methods to derive analytical approximations to model solutions in these cases.

Slow-growing astrocytomas were predicted to grow further into healthy tissue than faster, more malignant tumors, resulting in more edema over time. At the same time, changes in ICP were minimal in such tumors. This could explain why clinically, patients with grade I and II astrocytomas often exhibit symptoms only once their tumors have grown to a large extent. In contrast to those with astrocytomas of lower grade, patients with grade III and IV cancers typically present with acute symptoms. Model simulations revealed that for such faster growing tumors, the compensatory mechanisms of the brain are exhausted, inducing a sharp rise in ICP, especially near the tumor boundary. We remark that, as a first step towards a better understanding of tumor-induced edema and elevated ICP, we made the simplifying assumption of prescribing tumor growth rather than explicitly modeling the tumor itself. In future work, we plan to relax this assumption. Model validation – and extension to higher spatial dimensions – will be possible once increased clinical data in the form of SWE-derived real-time estimates of brain and cancer tissue stiffness become available.

One limitation of our model was that faster growing tumors such as GBM could not invade far into the brain, due to the volume fraction of healthy tissue rapidly approaching one near the tumor boundary. We therefore extended the scope of our model to include the effect that tumor-derived proteases may have on ICP levels and tumor invasion. Model simulations revealed that protease secretion was essential for increased tumor invasiveness, and may also be a potential mechanism underlying peritumoral edema onset.

Although simple, our model captures the biomechanical response of healthy brain tissue, in terms of changes in ICP and edema, to a growing tumor. Key differences in edema and pressure profiles were predicted that corresponded to tumors by grade. Thus, this model represents an important first step towards understanding the mechanisms that underlie ICP onset caused by brain cancer. We look forward to validating our model with clinical data as and when it becomes available. Once validated, such a model has the potential to improve brain cancer diagnostics via quantification of the (currently theoretical) notion of the Langfitt curve.

## AUTHOR CONTRIBUTIONS

ICS: performed research; analyzed data; wrote the manuscript.
MNJM: contributed analytic tools; wrote the manuscript.
HMB: designed research; contributed analytic tools; wrote the manuscript.
HVJ: designed research; contributed analytic tools; wrote the manuscript

## ACKNOWLEDGMENTS

This work was supported by the Simons Collaboration Grant for Mathematicians 280544 to HVJ.

